# Implementation and Evaluation of Support Vector Machine-Based Models for Cancer Detection Using Multi-Omic Data: A Systematic Review

**DOI:** 10.1101/2025.07.10.664049

**Authors:** Zhina Mohamadi, Erfan Abtahi, Zahra sadat Shayegh, Mehrafrin Ataei Kachouei, Amin Fakhar, Mohammad Mahdi Shirani, Mohammadhosein Malekian, Amir Zinatshoar, Mahdi Biglari, Fatemeh Rezaei, Armin Zarinkhat, Rozhina Mohammadi

## Abstract

**Introduction:** Cancer is a major source of mortality and morbidity all over the world that has caused more than 19 million new cases and nearly 10 million deaths in 2020. Although there are so many advances in cancer diagnosis, previous methods such as imaging and serum biomarkers more often lack the necessary sensitivity and specificity, particularly for early-stage detection.

However, most of the studies depend on internal validation that increases concerns about the generalizability of these findings. To improve the dependability of SVM applications in clinical fields, the review emphasizes the necessity of external validation and established techniques.

Due to all the things mixing AI with omics technology suggests a hopeful way to improve cancer detection, that could end up in better results and more affordable medical treatments.

**Method:** This systematic review was conducted using the PRISMA2020 principles and registered on The Open Science Framework. A comprehensive search of several databases was conducted, including PubMed/MEDLINE, Scopus, Google Scholar, and Web Of Science. Data was screened using RAYYAN.ai, which uses artificial intelligence methods to help with decision-making and screening. All original English-language studies that employed SVM to build a model for diagnosing a type human malignancy were included. The full text of the articles was extracted, and the quality of the articles and risk of bias were assessed using the PROBAST tool.

**Result:** A total of 104 studies were identified, of which 99 articles have been included after 5 were excluded because full text was unavailable. The studies covered various types of omics, such as proteomics (41 studies), transcriptomics (30 studies), genomics (19 studies), metabolomics (11 studies), epigenomics (4 studies), radiomics (2 studies), immunomics (1 study), and multi-omics (8 studies). 63 studies were internally validated, and 29 were externally validated; however, 2 studies were both internally and externally validated.

**Conclusion:** The review of 99 studies on Support Vector Machine-based models highlights their potential in improving cancer diagnosis. The study emphasizes the importance of proteomics studies in understanding tumor biology and developing effective diagnostic methods. However, concerns about their generalizability and trustworthiness in medical settings persist.

## Introduction

Cancer remains one of the leading causes of morbidity and mortality worldwide, with international data showing a global cancer-related death rate exceeding 21% as of 2012. In that year, 14.1 million new cancer cases were diagnosed and 8.2 million cancer-related deaths were recorded (1). The rise to over 19 million new cases and nearly 10 million deaths by 2020 emphasizes the ongoing need for effective prevention, early detection, and treatment strategies (2). Lung, breast, prostate, and colorectal cancers are among the most commonly diagnosed, accounting for almost half of all cancer-related deaths, with lung cancer being the most lethal among them (1, 3). These data highlight the critical need for more accurate and accessible methods of cancer detection to reduce mortality rates. While risk factors vary by cancer type, common contributors include lifestyle, genetic susceptibility, and environmental exposures. Specific predisposing factors have also been identified, such as tobacco use in lung cancer (4–7).

In spite of advancements in diagnostic tools, including biomarker identification and improved prognostic techniques, different cancer types continue to cause high mortality rates due to lack of detection accuracy and high diagnostic costs (8). Traditional methods such as imaging, biopsies, and serum biomarkers often do not have sufficient sensitivity and specificity, particularly in early-stage cancers. This can result in false positives, leading to unnecessary treatments, and false negatives, which delay critical interventions. The limitations of these approaches and the increment in cancer statistics clearly indicate the need for more precise, reliable, and cost-effective detection techniques with clinical applicability (9).

Omics technologies, which include transcriptomics, metabolomics, proteomics, and genomics, yield significant insights into the molecular causes of diseases like cancer and are modern approaches for biomarker exploration (3). These platforms enable the analysis of biological information at the levels of genes, RNAs, proteins, and metabolites, providing powerful tools for early diagnosis and prognosis by identifying cancer-specific molecular alterations (10). Each technology focuses on distinct molecular features: genomics investigates mutations, copy number variations, and structural alterations; proteomics examines protein expression profiles relevant to tumorigenesis; transcriptomics measures gene expression changes; and metabolomics explores disruptions in metabolic pathways (11). However, the volume and complexity of data generated by these platforms often exceed the capacity of traditional analytical methods, hence demanding the application of advanced computational tools (12).

Artificial intelligence (AI) can analyze such complex datasets by extracting structured and meaningful patterns through machine learning (ML) approaches (12). This computational strength is particularly beneficial in medical research, where conventional methods often lack accuracy and scalability. AI applications are also capable of analyzing radiological images, pathology slides, and omics datasets with high precision and efficiency. Support Vector Machines (SVMs) are highly effective in handling high-dimensional data among machine learning models, making them well-suited to process the complex outputs of omics technologies (2). Support Vector Machines (SVMs) have become well-known in cancer research as a widespread machine learning model, due to their remarkable ability to process high-dimensional biological data. Complex input data can be transformed into a higher-dimensional feature space employing a structure where various output classes are divided by an ideal hyperplane. Based on this characteristic and the large-margin classifier they utilize SVMs are extremely effective at differentiating between cancerous and non-cancerous patterns in omics data. This has made it possible for researchers to identify clinically significant patterns in transcriptomic, proteomic, and metabolomic profiles related to cancer detection, which has resulted in earlier and more accurate diagnoses.

By utilizing models such as SVM, researchers can find hidden trends in massive datasets, allowing for earlier and more accurate cancer detection. These models work by converting input data into a higher-dimensional feature space and then determining the best hyperplane for separating various classes. They use a large margin for classification, which enhances the detection of complex patterns with increased and more reliable accuracy (13, 14). This feature makes SVMs particularly effective for early-stage cancer diagnosis especially since omics platforms generate large and complex datasets (15, 16).

Although there are significant improvements at the intersection of omics technologies and computational approaches, including AI, there remains considerable untapped potential for translating these tools into clinical practice. Implementing such technologies in healthcare settings could contribute meaningfully toward reducing both cancer mortality and healthcare costs. This systematic review aims to compile and evaluate studies utilizing omics-based datasets analyzed through Support Vector Machines, with the goal of assessing their diagnostic performance, clinical potential, and contribution to early cancer detection. The findings aim to underscore the growing significance of AI-driven omics analysis in the future of oncology.

## Methods

This systematic review was conducted in accordance with the Preferred Reporting Items for Systematic Reviews and Meta Analyses (PRISMA2020) principles (17). The thorough checklist offered by PRISMA improves the planning and coordination of systematic reviews. Our systematic review has been registered on The Open Science Framework (OSF) (registration DOI: DOI 10.17605/OSF.IO/7VN5X)i.

## INFORMATION SOURCES and SEARCH STRATEGY

A detailed and comprehensive search of several databases from each database’s inception to August 4th 2024, was conducted. The databases included PubMed/MEDLINE, Scopus, Google Scholar and Web of Science. As seen in table 1, to achieve the goals of this topic a controlled vocabulary supplemented with keywords in each database was utilized.

**Table 1:** search strategy.

| DATABASE | SEARCH STRATEGY | RESULT |
| --- | --- | --- |
| PUBMED/MEDLINE | (("support vector machine"[Title/Abstract]) OR ("svm"[Title/Abstract])) OR ("Support Vector Machine"[Mesh]) AND (((("Neoplasms"[Title/Abstract]) OR ("Neoplasms"[Mesh])) OR ("carcinoma"[Title/Abstract]) OR ("carcinomas"[Title/Abstract])) OR ("Carcinoma"[Mesh])) OR ("sarcoma"[Title/Abstract]) OR ("sarcomas"[Title/Abstract])) OR ("Sarcoma"[Mesh]) AND (((((((((((((((((((("Metabolomics"[Title/Abstract]) OR ("metabolic profiling"[Title/Abstract])) OR ("metabolic fingerprinting"[Title/Abstract])) OR ("metabolic phenotyping"[Title/Abstract])) OR ("metabolome analysis"[Title/Abstract])) OR (epigenetic[Title/Abstract])) OR (epigenetics[Title/Abstract])) OR (epigenomics[Title/Abstract])) OR ("epigenetic profiling"[Title/Abstract])) OR ("epigenetic analysis"[Title/Abstract])) OR ("epigenetic modifications"[Title/Abstract])) OR ("epigenetic mechanisms"[Title/Abstract])) OR (proteomics[Title/Abstract])) OR ("protein analysis"[Title/Abstract])) OR ("protein profiling"[Title/Abstract])) OR ("Proteome research"[Title/Abstract])) OR ("proteome analysis"[Title/Abstract])) OR ("transcriptomics"[Title/Abstract])) OR ("gene expression profiling"[Title/Abstract])) OR | 677 |
|  | ("RNA sequencing"[Title/Abstract])) OR ("RNA-seq"[Title/Abstract])) OR ("transcriptome profiling"[Title/Abstract])) OR (genomics[Title/Abstract])) OR ("genome sequencing"[Title/Abstract])) OR ("genetic analysis"[Title/Abstract])) OR ("genome mapping"[Title/Abstract])) OR ("genomic profiling"[Title/Abstract])) OR ("genetic profiling"[Title/Abstract])) OR (Biomarker[Title/Abstract])) OR (biomarkers[Title/Abstract])) OR (gene[Title/Abstract])) OR (NGS[Title/Abstract])) OR ("Next generation sequencing"[Title/Abstract])) OR ("Micro RNA"[Title/Abstract])) OR (miRNA[Title/Abstract])) OR (protein[Title/Abstract])) OR ("Metabolomics"[Mesh])) OR ("Epigenomics"[Mesh])) OR ("Proteomics"[Mesh])) OR ("Gene Expression Profiling"[Mesh])) |  |
| WEB OF SCIENCE | (TS=(Metabolomics) OR TS=(metabolic profiling) OR TS=(metabolic fingerprinting) OR TS=(metabolic phenotyping) OR TS=(metabolome analysis) OR TS=(epigenetic) OR TS=(epigenetics) OR TS=(epigenomics) OR TS= ("epigenetic profiling") OR TS=("epigenetic analysis") OR TS=("epigenetic modifications") OR TS=("epigenetic mechanisms") OR TS=(proteomics) OR TS=("protein analysis") OR TS=("protein profiling") OR TS=("Proteome research") OR TS=("proteome analysis") OR TS=("transcriptomics") OR TS=("gene expression profiling") OR TS=("RNA sequencing") OR TS=("RNA-seq") OR TS=("transcriptome profiling") OR TS=(genomics) OR TS=("genome sequencing") OR TS=("genetic analysis") OR TS= ("genome mapping") OR TS=("genomic profiling") OR TS=("genetic profiling") OR TS= (Biomarker) OR TS=(biomarkers) OR TS=(gene) OR TS=(NGS) OR TS=("Next generation sequencing") OR TS= ("Micro RNA") OR TS=(miRNA) OR TS=(protein) OR TS=(genotype) OR TS=(phenotype)) AND (TS=(Support vector machine) OR TS=(SVM)) AND (TS=(Neoplasm) OR TS=(Neoplasms) OR TS=("carcinoma") OR TS=("carcinomas") OR TS=("sarcoma") OR TS=("sarcomas") OR TS=(malignant) OR TS=(malignancy) OR TS=(malignancies)) AND (TS=(Detection) OR TS=(diagnosis) OR TS=(diagnostic) OR TS= (detector) OR TS=(discovery)) | 642 |
| SCOPUS | (support vector machine OR SVM)<br>AND (Cancer OR Cancers OR tumor OR tumors OR malignancy OR malignancies OR Malignant OR carcinoma OR carcinomas OR sarcoma OR sarcomas OR neoplasm OR neoplasms OR "malignant growth") AND (metabolomics OR "metabolic profiling" OR "metabolic fingerprinting" OR "metabolic phenotyping" OR "metabolome analysis" OR epigenomics OR epigenetics OR "epigenetic profiling" OR "epigenetic analysis" OR "epigenetic modifications" OR "epigenetic mechanisms" OR proteomics OR "protein analysis" OR "protein profiling" OR "proteome research" OR "proteome analysis" OR "transcriptomics" OR "gene expression profiling" OR "transcriptome analysis" OR "RNA sequencing" OR "RNA-seq" OR "transcriptome profiling" OR genomics OR "genome sequencing" OR "genetic analysis" OR "genome mapping" OR "genomic profiling" OR "genetic profiling" OR Biomarker OR biomarkers OR gene OR NGS OR "Next generation sequencing" OR "Micro RNA" OR miRNA OR protein OR genotype OR phenotype)<br>AND (Detection OR Diagnosis OR Diagnostic OR Detector OR Discovery) | 2149 |
| GOOGLE SCHOLAR | allintitle: "support vector machine" "metabolomics" OR "phenotype" OR "genotype" OR "metabolic profiling" OR "metabolic fingerprinting" OR "metabolic phenotyping" OR "metabolome analysis" OR "epigenetic" OR "epigenetics" OR "epigenomics" OR "epigenetic profiling" OR "epigenetic analysis" OR "epigenetic modifications" OR "epigenetic mechanisms" OR "proteomics" OR "protein analysis" OR "protein profiling" OR "Proteome research" OR "proteome analysis" OR "transcriptomics" OR "gene expression profiling" OR "RNA sequencing" OR "RNA-seq" OR "transcriptome profiling" OR "genomics" OR "genome sequencing" OR "genetic analysis" OR "genome mapping" OR "genomic profiling" OR "genetic profiling" OR "Biomarker" OR "biomarkers" OR "gene" OR "NGS" OR "Next generation sequencing" OR "Micro RNA" OR "miRNA" OR "protein" "cancer" | 14 |
| Total |  | 3482 |

## DATA SCREENING AND ELIGIBILITY CRITERIA

We used RAYYAN.ai to screen the search results (18). RAYYAN uses different artificial intelligence methods to do a great help with decision making and screening in systematic and literature reviews. Titles and abstracts from 3482 articles retrieved from our search strategy were blindly and independently screened by seven reviewers (E.A, A.F, Zh.M, Z,Sh, M.B, F.R, A.Z). RAYYAN ai was employed to eliminate the duplicated records. The conflicts were resolved by another independent reviewer (A.Zkh).

All of the original English-language studies that employed SVM to build a model for diagnosing a type human malignancy were included in this study (table 2). During the initial screening phase, the inclusion criteria were only examined at the level of the articles’ titles and abstracts. However, in the subsequent phase, we screened the full text of included articles. Included articles whose full text was not available were excluded from the review.

**Table 2:** step by step checklist to include an article. Failure to meet the criteria of each step led to exclusion.

|  |
| --- |
| 1. A Journal article |
| 2. An English language article |
| 3. A human malignancy |
| 4. Developing an diagnostic model |
| 5. Application of SVM in model development |
| 6. Utilizing an omic to build the model |
| 7. Article full text available |

## DATA EXTRACTION AND QUALITY ASSESSMENT

In the subsequent phase of screening, we used the included articles’ full text to extract the following data: Author, Title, Year, Country, aim of study, population, Cancer type, Use of SVM in the model, Type of validation, Type of omic, Factor(s) used for diagnosis, Method, Outcome, Conclusion and Quality of evidence.

The type of validation in models that used a population other than the sample population specified in the data extraction process to confirm the application of the model was considered external validation. How robust the results are when applied to the training dataset is the main focus of internal validation, while external validation results can be generalized to the reference population (19).

To assess the quality of the included articles and risk of bias, four assessors (E. A, M. A, M, Sh, Z. Sh), evaluated each study separately based on the PROBAST tool. The tool comprises four domains: Participants, predictors, outcome and analysis. These domains contain a total of 20 signaling questions to facilitate structured judgment of risk of bias, which was defined to occur when shortcomings in study design, conduct, or analysis led to systematically distorted estimates of model predictive performance (20).

## Result

### Study Selection

Our comprehensive search in the four aforementioned databases yielded a total number of 104 studies though we were unable to obtain the full text of five papers.

From a total of 99 articles, 63 presented internal validation,29 presented external validation and 2 presented both internal and external validation (figure 1).

**Figure 1:**
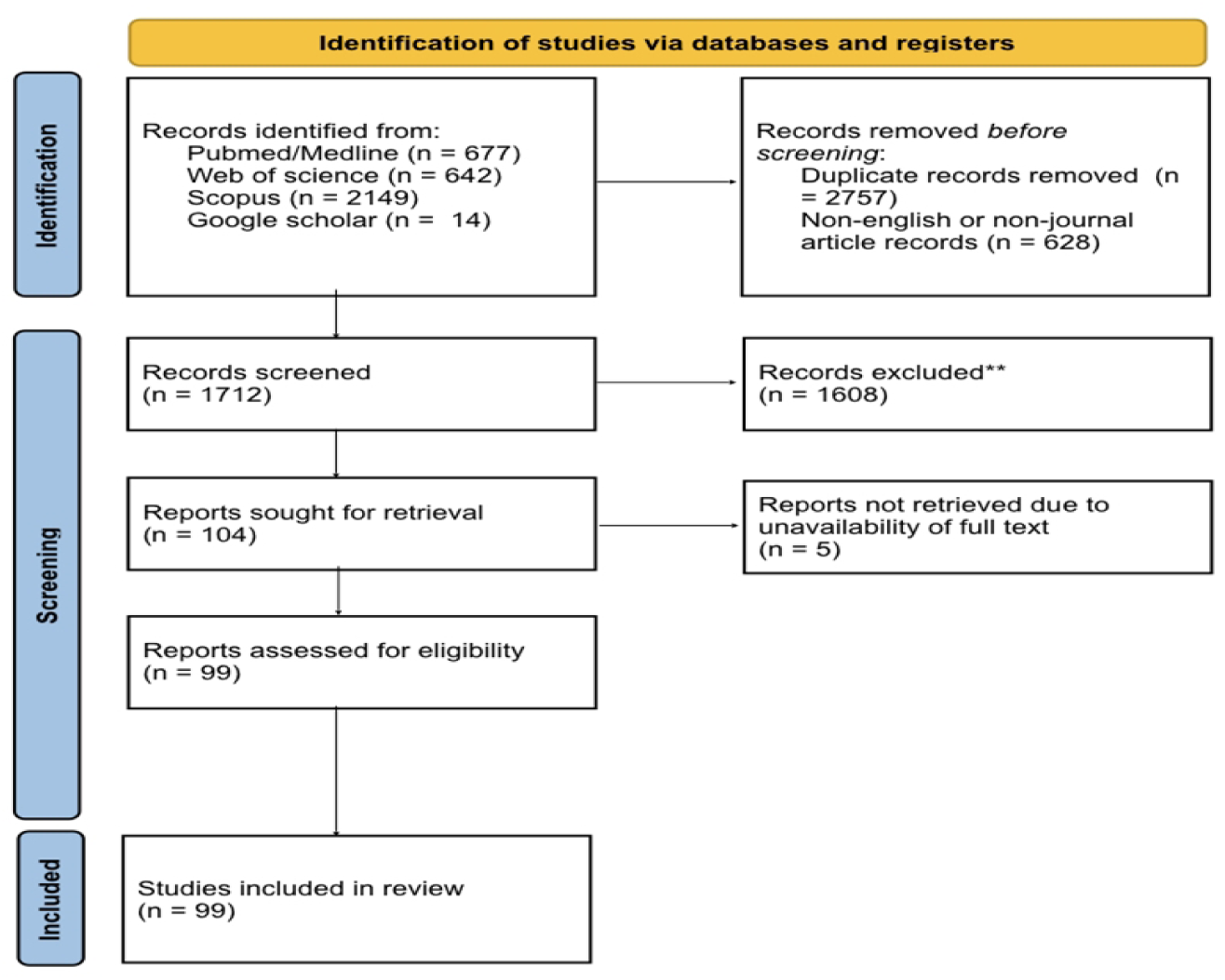
PRISMA 2020 flow diagram for new systematic reviews which included searches of databases

### Study Characteristics

We reviewed a range of studies including clinical trials and observational studies that were published between 2005 and 2024. They were performed in different countries with most of them occurring in USA and China (table 3).

**Table 3:**
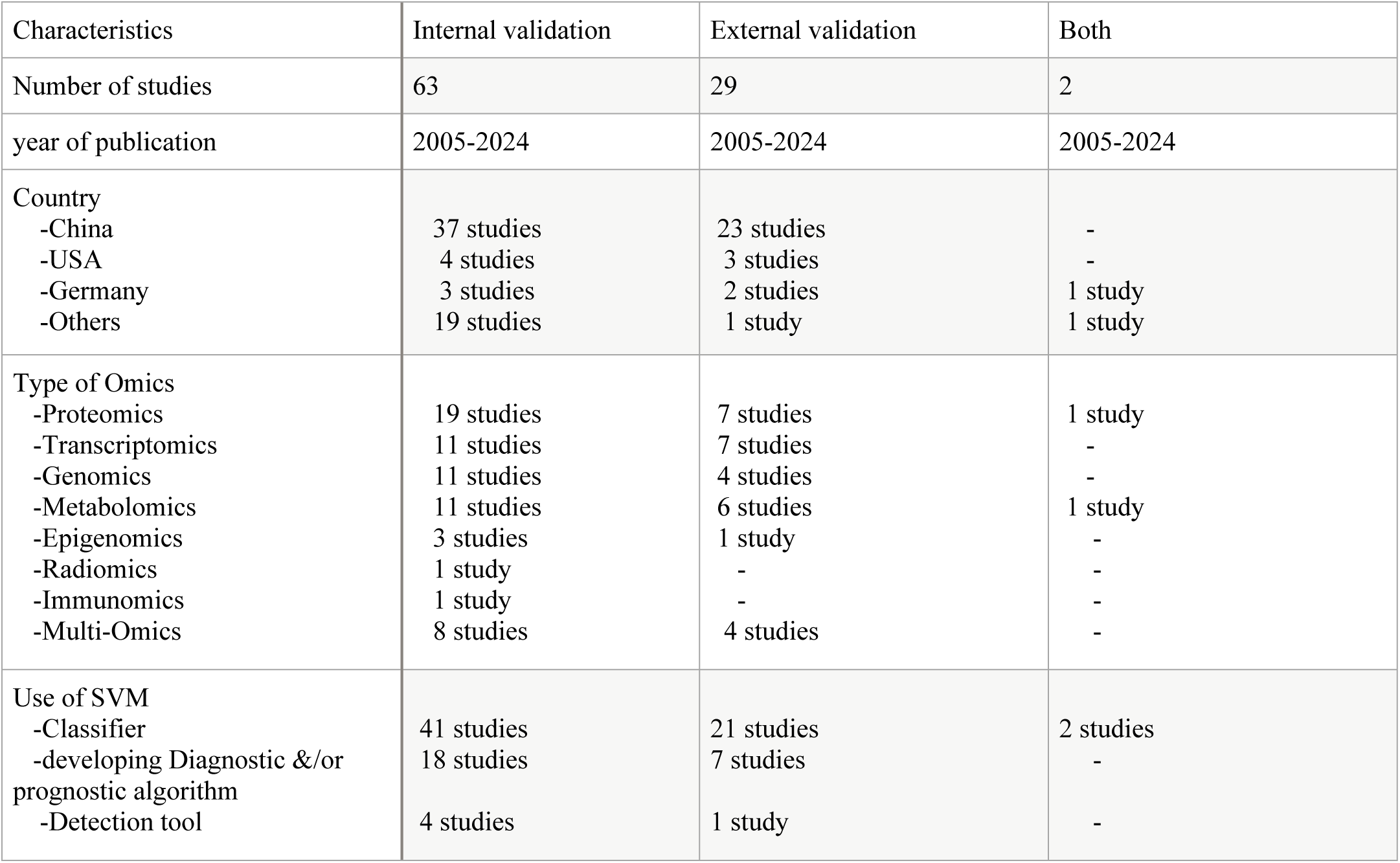
Studies which investigate internal validation, external validation, and both.

A total of 99 studies were included in this review, of which 63 utilized internal validation, 29 applied external validation, and only 2 employed both. Internal and external validation are clearly different. Whilst external validation measures model performance on several datasets, internal validation uses the same dataset or a portion of it. The absence of external validation restricts generalizability, as we demonstrated in our previous study, highlights the necessity of strong validation to guarantee clinical application (129). Type of validation in the remaining studies was not determined. The publication years spanned from 2005 to 2024 across all validation categories. Most studies originated from China, particularly in the internal validation group (37 out of 63), suggesting regional dominance in SVM-based omics research. In terms of omics data types, proteomics was the most frequently used across all validation strategies, followed by transcriptomics, genomics, and metabolomics. Multi-omics approaches were also employed in a modest number of studies, especially in internally validated research. Regarding SVM application, the majority of studies used it as a classifier (particularly in internally validated research), while fewer aimed to develop diagnostic or prognostic models.

### Risk of bias

The risk of bias was assessed for all of 99 articles using PROBAST tool, and most of them had moderate risk of bias with a much smaller group having high risk of bias (table 4).

**Table 4:** The risk of bias was assessed for all of 99 articles using PROBAST tool.

| Risk of Bias based on PROBAST | Number of Articles | Percentage (%) |
| --- | --- | --- |
| Low | 34 | 34.3 |
| Moderate | 47 | 47.7 |
| High | 15 | 15.1 |

The risk of bias assessment using the PROBAST tool showed that approximately one-third of the included studies (34.3%) were classified as having a low risk of bias, while nearly half (47.7%) demonstrated a moderate risk. Only 15.1% of studies were deemed to have a high risk of bias.

### Findings

In the reviewed studies, it was shown that based on the application of SVM, employed omics were useful for either diagnosis, prognosis, or classification of different types of cancers with mostly high sensitivity and specificity with some of them even reaching an AUC=1.00 and 100% sensitivity. One of which was W. Chen’s 2023 study in USA, which used CT scans to detect early changes on computed tomography (CT) images associated with pancreatic ductal adenocarcinoma (PDAC) based on quantitative imaging features (QIFs) for patients with and without chronic pancreatitis (CP). The pancreas was automatically segmented, and 111 QIFs were extracted. Conditional support vector machine (SVM) algorithms were created individually for patients with and without CP. That resulted in the accuracy measures for non-CP patients were 94%-95%, with AUC measurements ranging from 0.98 to 0.99. For patients with CP, the accuracy was 100%, with an AUC of 1.00. As a result, it found that QIFs can accurately predict PDAC for both non-CP and CP patients on CT imaging, indicating prospective biomarkers for early identification of pancreatic representing promising biomarkers for early detection of pancreatic cancer. The study was effective in demonstrating the predictive capability of the algorithms.

In table 5, some of the papers which have low risk of bias and used external validation at the same time are listed with relative outcomes. Also, the following table represents the range of cancer types that were covered in the reviewed literature (table 5). The diagnostic outcomes reveal that support vector machine (SVM)-based models show promising performance across various cancer types when applied to omics datasets. We categorized SVM diagnostic applications by cancer group.

**Table 5:**
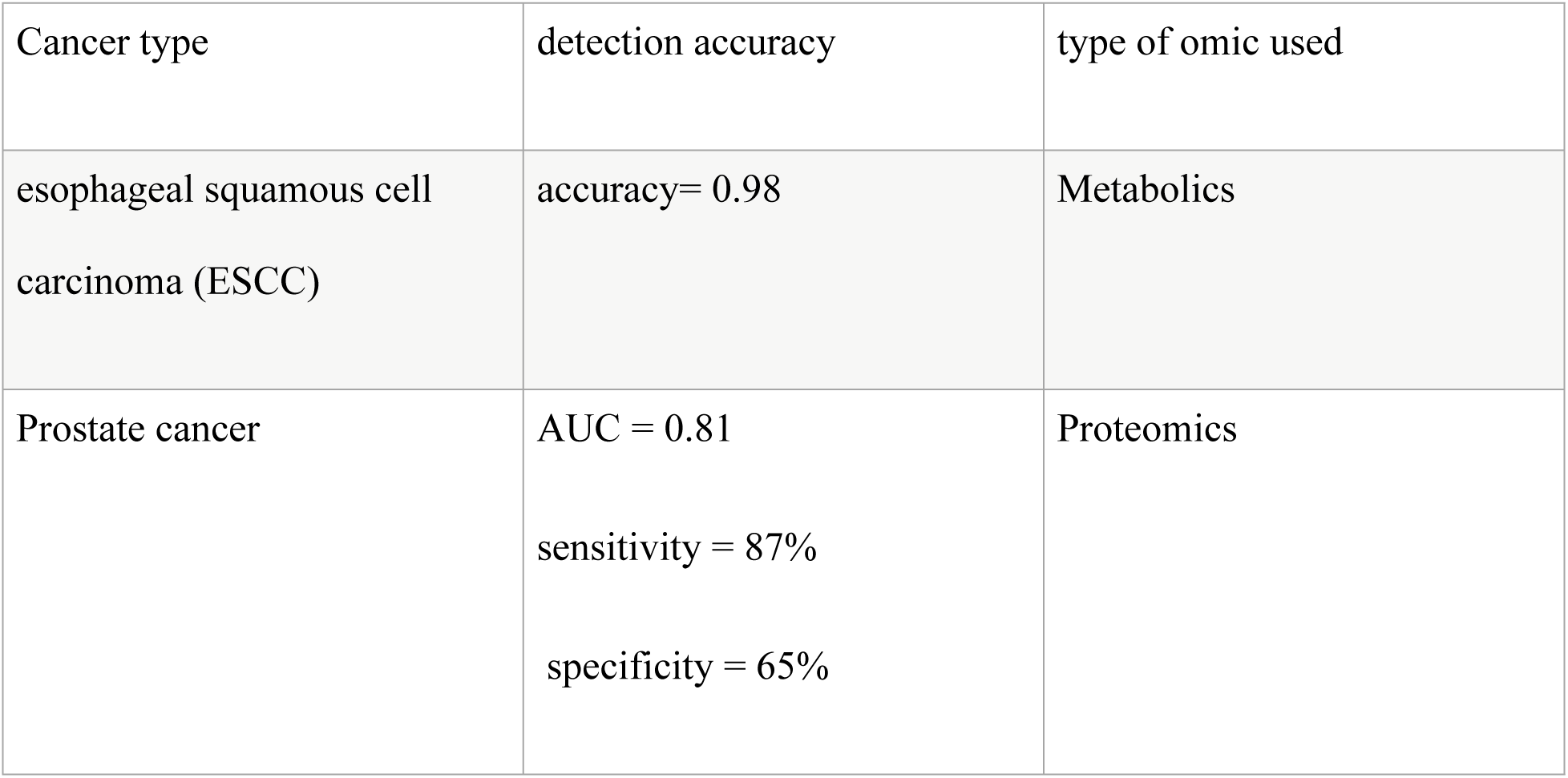
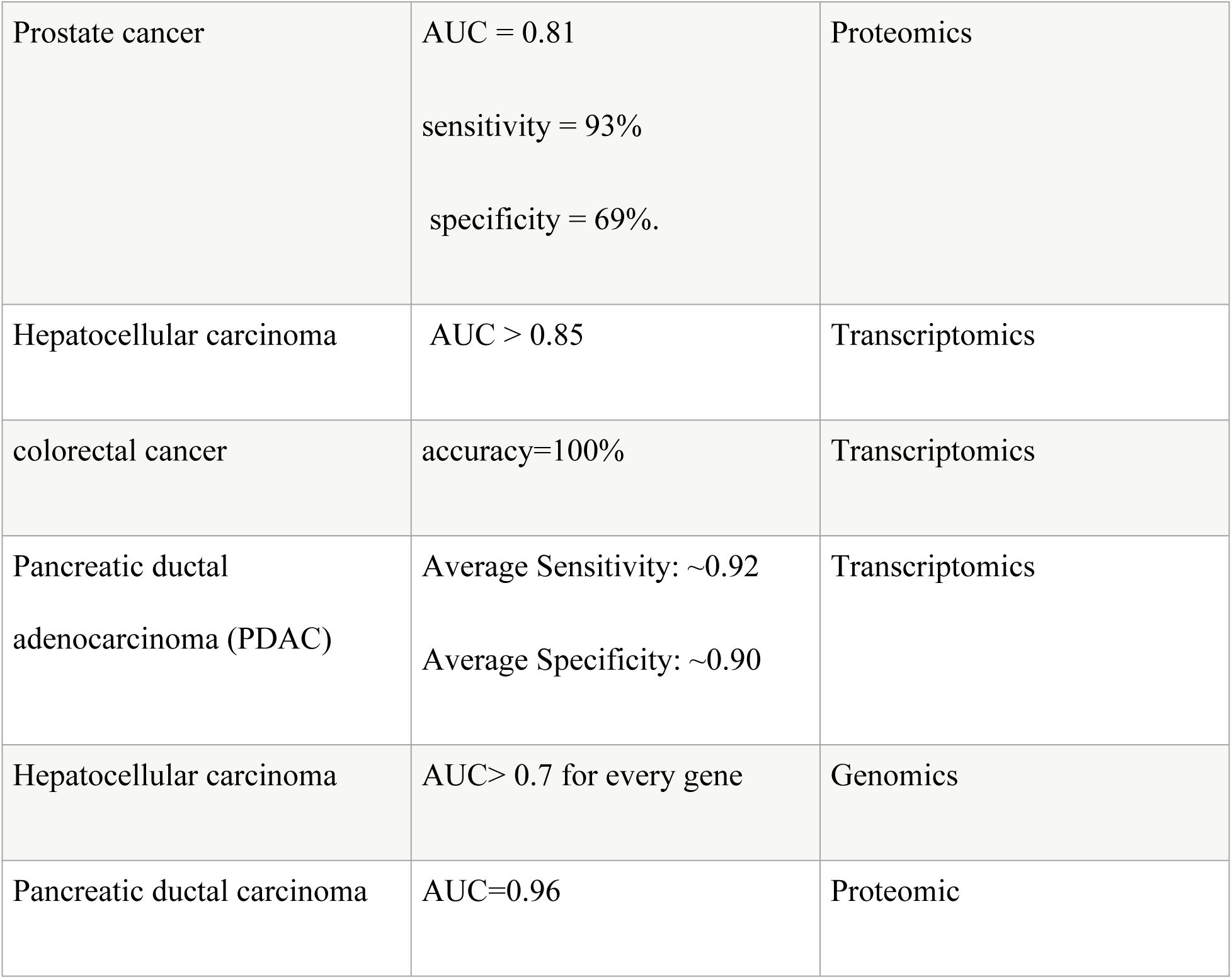
represents the range of cancer types that were covered in the reviewed literature.

| Cancer type | detection accuracy | type of omic used |
| --- | --- | --- |
| esophageal squamous cell carcinoma (ESCC) | accuracy= 0.98 | Metabolics |
| Prostate cancer | AUC = 0.81<br>sensitivity = 87%<br>specificity = 65% | Proteomics |
| Prostate cancer | AUC = 0.81<br>sensitivity = 93%<br>specificity = 69%. | Proteomics |
| Hepatocellular carcinoma | AUC > 0.85 | Transcriptomics |
| colorectal cancer | accuracy=100% | Transcriptomics |
| Pancreatic ductal<br>adenocarcinoma (PDAC) | Average Sensitivity: ~0.92<br>Average Specificity: ~0.90 | Transcriptomics |
| Hepatocellular carcinoma | AUC> 0.7 for every gene | Genomics |
| Pancreatic ductal carcinoma | AUC=0.96 | Proteomic |

Hematologic Malignancies:

Hodgkin’s Disease: A serum proteomic SVM model (CE-MS profiling) achieved 98% sensitivity and 97% specificity in leave-one-out cross-validation (AUC = 0.99).

Leukemia Subtypes: Transcriptomic SVM classifiers distinguished acute myeloid leukemia (AML) from acute lymphoblastic leukemia (ALL) with 100% accuracy in independent microarray cohorts

Gastrointestinal (GI) Malignancies:

Esophageal Squamous Cell Carcinoma: Plasma-metabolomics SVM yielded 98% accuracy (AUC = 0.98).

Colorectal Cancer: An integrated transcriptomic SVM achieved 100% accuracy in two external validation sets (AUC = 1.00).

Pancreatic Ductal Adenocarcinoma: Radiomic SVM models based on CT-derived quantitative imaging features distinguished PDAC from chronic pancreatitis with 94–100% accuracy (AUC = 0.98–1.00).

In hepatocellular carcinoma (HCC)Transcriptomic SVM-RFE Identified VIPR1 and co-expressed genes, achieving external-validation AUC > 0.85 and Genomic Signature SVM with HBV-related gene panels classified TCGA HCC versus controls with AUC = 1.00 and correctly classified 78/82 samples in independent cohorts.

Genitourinary (GU) Malignancies:

Prostate Cancer: Urinary proteomic SVM classifiers (CE-MS peptides) reported external-validation AUC = 0.81, sensitivity = 93%, specificity = 69%.

Renal Cell Carcinoma: Serum lipidomic SVM panels achieved training AUC = 0.95 and test AUC = 0.82.

Other Solid Tumors:

Breast Cancer: Serum peptidome SVM models attained external-validation sensitivity and specificity > 91%.

Nasopharyngeal Carcinoma: Hyperspectral-imaging SVM classifier achieved 99.2% accuracy (AUC = 0.98).

Thyroid Nodules: Transcriptomic SVM distinguished benign from malignant nodules with 95% sensitivity and 92% specificity.

Across all groups, SVM models demonstrated high diagnostic performance, with most reporting sensitivities and specificities ≥ 90% and several perfect classifications (AUC = 1.00).

These findings suggest that omics-based SVM models—particularly those utilizing transcriptomic and proteomic data—can offer high diagnostic accuracy and clinical potential, though performance varies by cancer type and omics layer.

To provide a comprehensive overview of the current landscape of SVM-based cancer detection using omics data, we compiled the distribution of included studies based on cancer type, the type of omics data utilized, and the validation approach employed. Table 6 summarizes the frequency with which each omics platform (e.g., proteomics, transcriptomics, genomics, metabolomics, epigenomics, radiomics, immunomics, and multi-omics) has been applied across different cancer types. Additionally, it outlines the validation strategies used in these studies—internal, external, or both. Notably, gastrointestinal and genitourinary cancers were the most frequently studied categories across multiple omics layers. Multi-omics integration was employed in several cancer types, though still less frequently compared to single-omics approaches. Internal validation was more commonly reported than external validation, with only a limited number of studies conducting both. This distribution highlights both the diversity and imbalance in current SVM-based omics applications and suggests opportunities for expanding multi-omics strategies and rigorous external validation in future research.

**Table 6:** Summary of included studies categorized by cancer type, omics platform used (proteomics, transcriptomics, genomics, etc.), and type of model validation (internal, external, or both). Each number indicates the count of studies applying the respective omics platform or validation strategy to the specified cancer type. Multi-omics refers to studies combining two or more omics data types for model development.

| Cancer Type | Proteomics | Transcriptomics | Genomics | Metabolo-omics | Epigenomics | Radiomics | Immunomics | Multi-Omics | Internal Validation | External Validation | Both |
| --- | --- | --- | --- | --- | --- | --- | --- | --- | --- | --- | --- |
| Gastrointestinal tract | 6 | 5 | 5 | 3 | 3 | - | - | 2 | 20 | 5 | - |
| Genitourinary system | 6 | 2 | 1 | 6 | - | - | - | 1 | 10 | 5 | 2 |
| Hepatocellular | 4 | 1 | 3 | - | - | - | 1 | 1 | 6 | 5 | - |
| Pancreatic | 2 | 4 | 1 | - | - | 1 | - | 2 | 3 | 5 | - |
| Breast | 2 | 2 | 1 | 3 | - | - | - | 2 | 6 | 2 | - |
| Hematologic | 2 | - | 1 | - | - | - | - | 1 | 4 | - | - |
| Lung | 4 | 2 | - | - | 1 | - | - | 1 | 5 | 3 | - |
| Thyroid | 1 | 2 | - | - | - | - | - | - | 2 | 1 | - |
| CNS | - | - | 1 | 1 | - | - | - | 1 | 2 | 1 | - |
| Other | 1 | 1 | 2 | 4 | - | - | - | 2 | 4 | 2 | - |

As the above table demonstrates, proteomics is by far the most commonly used omics across multiple cancer research, and gastrointestinal tract neoplasia is the most frequently studied type of malignancy. Further, there appears to be a tentative pattern suggesting that studies using metabolomics and transcriptomics more frequently employed external validation than those using proteomics.

## Discussion

In this systematic review, we analyzed 99 studies on the outcome of support vector machine (SVM) algorithms when different omics data were inputted into the algorithm to determine a more efficient prognostic and diagnostic method in cancer detection. Our findings reveal valuable insight into the efficacy of these models across various cancer types and omics applications.

### Key Findings and Patterns

The internal validation process evaluates the model’s outcome based on the same data used in the development of the model. Cross-validation is implied to prevent reliance on specific input data. External validation, on the other hand, assesses the data not originally used in developing the model. This step is necessary for the evaluation of the model’s generalizability. We acquire insights about the reliability of the model’s performance in real-life situations by externally validating it (118). This review indicated a noticeable internal validation rate (63 out of 99), whereas only 29 studies included external validation, and 2 used both methods. This highlights the necessity of more externally validated research to prove these SVM-based models’ efficacy in clinical settings and their ability to be generally used. R. Riley et al have done a systematic review emphasizing the importance of external validation. This review indicated that external validation shows the reliability of the model’s performance in different patient groups and clinical settings (119). Another study also highlights that if this step is overlooked, applying the model in real-life situations will have a considerable risk of inaccuracy (120).

The diversity of countries involved in a study can significantly affect the outcome of a systematic review. If a considerable number of studies are done in just a few countries, the conclusion can be biased and incapable of representing the global population (121, 122). Given this information, there is a geographic focus on the amount of research done in China and the USA, which may compromise the applicability of our findings in different populations.

Proteomics addresses the extensive examination of proteins. This field of study provides the knowledge for the identification and quantification of protein expression levels, as well as the analysis of the variables post-translation and protein interactions. Such information has proved to be crucial in understanding tumor biology (123, 124). This omics technology emerged as the most researched approach, followed by transcriptomics and genomics. This trend indicated the possibility of a correlation between proteomics and important insights into cancer biology, the development of diagnostic methods, and efficient prognosis.

Different types of cancer were analyzed in this study; notably, gastrointestinal tract malignancies were the most studied. Research indicates that the broader the scope of the studies, including a wider range of cancer types, the more generalizable they are. On the contrary, a restricted focus on specific types of cancer can lead to a more concentrated result, which is not the purpose of systematic review articles (125, 126).

### Expectations and Comparisons

Assessing the risk of bias (ROB) is essential for validating and applying the findings effectively. There are several tools available for ROB assessment, one of which is PROBAST (Prediction model Risk of Bias Assessment Tool). This tool has become well-known for its structured framework (including 20 signaling questions that support four fields of bias evaluation), enabling PROBAST to be more specifically designed for studies involving predicting models and thoroughly evaluating potential biases that could twist the model’s performance (20). Our assessment of bias using the PROBAST tool revealed a 47.7% moderate risk of bias, while 15.1% were classified as having a high risk. This requires a more accurate methodology for further research before clinical applications.

One remarkable category of highly effective machine-learning algorithms is Support Vector Machines (SVM), which are vastly utilized in medical diagnostics due to their impressive accuracy in classifying complex data. Recent studies suggest that SVM can be an advantageous tool in different medical fields. For instance, a study on the diagnosis of breast cancer indicated an accuracy rate of 99.25%, along with sensitivity and specificity rates of 98.96% and 100%.

This highlights SVM’s accuracy in identifying cancerous tissues (127). The sensitivity and specificity of SVM models in our findings were measured high, and this result aligned with our expectations. Several studies even reported an area under the curve (AUC) of 1.00, reaching 100% sensitivity, which signifies the SVM algorithm’s high potential as a diagnostic tool in the medical setting. For instance, W. Chen’s 2023 study demonstrates the capability of quantitative imaging features (QIFs) to predict pancreatic ductal adenocarcinoma (PDAC) with 100% accuracy, supporting the theory of using multi-omics data analysis of SVM models to enhance diagnostic accuracy (27).

When comparing our findings with past research, we see a growing use of machine learning techniques, especially SVM, in oncology. For instance, a study indicated efficient outcomes in cancer genomics classification when using SVM, which led to identifying biomarkers, novel drug targets, and enhanced insights into cancer driver genes (128). SVMs’ capability to manage high-dimensional data makes them invaluable tools, especially for analyzing gene expression profiles and various omics datasets (127). However, our findings extend the science gap by emphasizing the critical need for more externally validated research to apply the promising SVM models in clinical settings.

### Unexpected Findings and Alternative Explanations

One unexpected finding was the relatively high rate of studies labeled moderate to high risk of bias, which raises concerns about the rigor of many studies’ methodologies. Despite the high risk of bias, one factor that indicates favorable accuracy metrics is Internal over external validation dominance due to its unrealistic reflection of results.

The observed pattern might also be affected by research priorities in a region or the unavailability of funding in other areas, specifically in China (60 studies) and the USA (7 studies). Furthermore, the dominance of proteomics research (27 studies) might be related to advancements in this field and not necessarily the superiority of this approach.

### Implications and Practical Applications

Our findings show the significant potential of SVM models to enhance detection accuracy through multi-omic data. High sensitivity and specificity of these models in several studies were reported, particularly in early-stage cancer detection, where timely intervention can greatly impact the patient’s healing process. However, the need for further validation highlights the importance of establishing protocols and standard methods to ensure the reliability of these models in diverse clinical settings.

## Limitations

### Heterogeneity Across the Dataset

A critical observation is the high heterogeneity across the studies in several aspects: cancer types, omics modalities, model validation strategies, dataset sizes, and performance metrics. This variability makes direct comparison between studies difficult and impairs the possibility of meta-analytic synthesis. For instance, performance measures such as AUC, accuracy, sensitivity, and specificity were not reported consistently, and when reported, often used different thresholds or calculation methods. This heterogeneity is further exacerbated by the diversity in feature selection techniques and SVM parameter tuning, leading to outcome variability that is methodological rather than biological. Addressing this heterogeneity through standardization and stratified analysis is crucial for deriving robust and generalizable insights.

It is essential to acknowledge the limitations of our research as some of them are pointed out in the results sections as findings of the data extraction table. The diversity of study design and methodology in 99 articles can create potential confounding variables that could challenge our ability to classify and compare results accurately. Also, the high performance of specific countries in this field, as mentioned in the findings of the results, can alter the generalizability of our findings worldwide.

Another limitation is the high heterogeneity across the studies in several aspects: cancer types, omics modalities, model validation strategies, dataset sizes, and performance metrics. This variability makes direct comparison between studies difficult and impairs the possibility of meta-analytic synthesis for our systematic review. For instance, performance measures such as AUC, accuracy, sensitivity, and specificity were not reported consistently, and when reported, often used different thresholds or calculation methods. Addressing this heterogeneity through standardization and stratified analysis is crucial for deriving robust and generalizable insights.

Therefore, while standardization and stratified analyses would enhance comparability, the existing diversity does not substantially compromise the overall interpretability or utility of the findings.

The use of proteomics was associated with lower quality of evidence in the included studies as mentioned in the results section. This may be due to the broader availability of public transcriptomic datasets (e.g., TCGA, GEO) compared to proteomics, which often requires customized experimental data. This distinction is important as it could indicate that certain omics types are more conducive to model generalization and reproducibility due to greater data accessibility.

The risk of bias observed in the collected studies can raise great concern about the reliability of the predicted reports on the performance of SVM models. Moreover, the lack of rigor in external validation in most of the reviewed research limits our ability to make a final deduction regarding the clinical applications of these models.

Further, Despite the theoretical advantage of integrating multiple omics layers to capture the complexity of cancer biology, relatively few studies employed a true multi-omics approach.

Most studies relied on a single omics layer, and only a minority applied combined data from two or more omics sources.

### Recommendations for Future Research

Increasing the efficacy and reliability of SVM models is one of the underlying purposes of our study. To enhance this matter, we gathered a few recommendations:

1. Standardizing the Methodologies: Upcoming research should develop standard guidelines for gathering and analysis of data. This would simplify and facilitate comparing different methods and improve the reliability of research.
2. Focusing on External Validation: A comprehensive emphasis on external validation in upcoming research ensures more reliable results concerning the use of SVM models across various medical environments.
3. Exploring Underrepresented Cancer Types: Investigating a larger range of cancer types, especially those currently less engaged in this field of research could result in a deeper insight into the application of SVM models.
4. Integrating Multi-Omics Approaches: Future research should focus on further exploring all various omics fields to predict the function of SVM models in every aspect necessary.
5. Longitudinal Studies for Dynamic Monitoring: Extended methodologies can provide long-term insight into tumor development. The effects of early cancer detection by SVM models and treatment responses could also be predicted. These methods may enhance personalized medical strategies worldwide.
6. Including Clinical and Demographic Variables: Adding health factors and demographic information alongside omics data could enhance predictive accuracy.
7. Employing Advanced Machine Learning Techniques: Exploring additional advanced machine learning techniques could offer improved management and classification of complex, multi-dimensional data or at least provide an alternative insight into employing SVM models in this field of study.

In conclusion, while our systematic review offers helpful insight into the application of SVM for cancer detection, it also highlights blind spots that leave space for improvement. These recommendations will contribute to developing more efficient diagnostic tools and ultimately early-stage cancer detection and better patient outcomes in oncology.

## Conclusion

This systematic review of 99 studies demonstrates the remarkable diagnostic and prognostic potential of Support Vector Machines (SVMs) in cancer diagnosis through the study of omics data. Despite the models’ consistently high sensitivity and specificity—particularly for early-stage cancer—their over-reliance on internal validation limits their practical application. There are questions regarding the generalizability and clinical usefulness of these models because only a small percentage of research underwent external validation.

Despite being commonly linked to lower-quality evidence, proteomics has become the most used omics method. Transcriptomic-based research, on the other hand, seemed to gain from enhanced information accessibility, which might lead to more repeatable results. The conclusions’ potential to be applied globally may be restricted by the studies’ notable regional concentration in a few numbers of nations, including China and the USA.

Synthesis and comparison are made more difficult by the high degree of variation across research, for instance in terms of cancer types, omics layers, validation procedures, dataset sizes, and performance indicators. Comprehensive conclusions are hampered by inconsistent reporting and underutilization of multi-omics integration, which emphasizes the importance of standardized procedures.

Future studies should concentrate on external validation, standardizing methodological approaches, investigating underrepresented malignancies, and integrating multi-omics data in order to address these gaps. Predictive accuracy and relevance would also be improved by adopting longitudinal designs and incorporating clinical and demographic parameters. In conclusion, converting SVM-based omics models into clinically valid instruments for early cancer detection and better patient outcomes demands increased scientific accuracy and decreased bias.

## Declarations

### Registration and protocol

The review was not registered. A protocol was not prepared.

## Abbreviations

Not applicable.

## Ethics approval and consent to participate

This study does not involve human participants, animal subjects, or patient data. Therefore, ethics approval and consent to participate are not required.

## Consent for publication

The results/data/figures in this manuscript have not been published elsewhere, nor are they under consideration by another publisher.

## Availability of data and materials

This study is a systematic review and does not involve the generation of new datasets. All data analyzed in this study were obtained from previously published sources, which are cited in the manuscript.

## Authors’ contributions

Zh.M, E.A, Z.SH, and A.F wrote the main manuscript.

M.A, M.Sh, M.M, A.Z, M.B, R.M, F.R contributed in data collection, screening, extraction and quality assessment.

A.Zkh prepared the figures.

All authors reviewed the article.

## Acknowledgements

Not applicable.

## Footnotes

i OSF [Internet]. Available from: https://osf.io/

